# Evaluating the accuracy of genomic prediction of growth and wood traits in two *Eucalyptus* species and their F_1_ hybrids

**DOI:** 10.1101/081281

**Authors:** Biyue Tan, Dario Grattapaglia, Gustavo Salgado Martins, Karina Zamprogno Ferreira, Björn Sundberg, Pär K. Ingvarsson

## Abstract

**Background:** Genomic prediction is a genomics assisted breeding methodology that can increase genetic gains by accelerating the breeding cycle and potentially improving the accuracy of breeding values. In this study, we used 41,304 informative SNPs genotyped in a *Eucalyptus* breeding population involving 90 *E.grandis* and 78 *E.urophylla* parents and their 949 F_1_ hybrids to develop genomic prediction models for eight phenotypic traits - basic density and pulp yield, circumference at breast height and height and tree volume scored at age thee and six years. Based on different genomic prediction methods we assessed the impact of the composition and size of the training/validation sets and the number and genomic location of SNPs on the predictive ability (PA).

**Results:** Heritabilities estimated using the realized genomic relationship matrix (GRM) were considerably higher than estimates based on the expected pedigree, mainly due to inconsistencies in the expected pedigree that were readily corrected by the GRM. Moreover, GRM more precisely capture Mendelian sampling among related individuals, such that the genetic covariance was based on the actual proportion of the genome shared between individuals. PA improved considerably when increasing the size of the training set and by enhancing relatedness to the validation set. Prediction models trained on pure species parents could not predict well in F_1_ hybrids, indicating that model training has to be carried out in hybrid populations if one is to predict in hybrid selection candidates. The different genomic prediction methods provided similar results for all traits, therefore GBLUP or rrBLUP represents better compromises between computational time and prediction efficiency. Only slight improvement was observed in PA when more than 5,000 SNPs were used for all traits. Using SNPs in intergenic regions provided slightly better PA than using SNPs sampled exclusively in genic regions.

**Conclusions:** Effects of training set size and composition and number of SNPs used are the most important factors for model prediction rather than prediction method and the genomic location of SNPs. Furthermore, training the prediction model on pure parental species provide limited ability to predict traits in interspecific hybrids. Our results provide additional promising perspectives for the implementation of genomic prediction in *Eucalyptus* breeding programs.

## Background

*Eucalyptus* species and their hybrids are the most widely planted hardwoods in tropical, subtropical and temperate regions, due to their fast growth, short rotation, wide environmental adaptability and suitability for commercial pulp and paper production [1, 2]. Interspecific hybrids of *E.grandis* and *E.urophylla*, in particular, are generally superior to their parents in growth, wood quality and biotic and abiotic stresses resistance, by inheriting both the fast growth and good rooting abilities of *E.grandis* and the disease tolerance and wide adaptability of *E.urophylla* [3]. A conventional breeding cycle toward clonal selection in hybrid populations involves mating, progeny trial, a small-scale clonal trial and a second expanded clonal trial, that together typically take between 12 and 18 years [1, 4]. To accelerate the genetic gain per unit time, new methods that can help shorten the breeding cycles are greatly needed.

Genomic prediction or genomic selection (GS) is one of the most recent developments in genomics-assisted methods that are aimed at improving breeding efficiency and genetic gains. Genomic prediction provides a genome-wide paradigm for marker-assisted selection (MAS)[5, 6]. In GS all markers are fitted simultaneously in a model that relies on the principle of linkage disequilibrium (LD) to capture most of the relevant variation throughout the genome, whereas MAS focuses on discrete quantitative trait loci (QTLs) that had previously been detected, usually in underpowered experiments and thus leaving most of the variation unaccounted for [7]. GS are generally performed in three steps: (1) genotyping and phenotyping a ‘reference’ or ‘training population’ and developing genomic prediction models that allow for prediction of phenotypes from genotypes; (2) validation of the predictive models in a ‘validation population’, i.e. a set of individuals that did not participate in model training; (3) application of the models to predict the genomic estimated breeding values (GEBVs) of unphenotyped individuals which are then selected according to their GEBVs [6]. GS has been successfully implemented in the breeding of livestock [7, 8] and crops [9, 10] and several recent papers suggest that has great potential also in forest trees [11, 12].

The accuracy of genomic prediction models can vary depending on the statistical method employed. Several methods have been developed for GS, including ridge-regression best linear unbiased prediction (rrBLUP), genomic best linear unbiased prediction (GBLUP), BayesA, BayesB, Bayesian LASSO, BayesR and reproducing kernel Hilbert space (RKHS) regression [7, 13]. These methods vary in the assumptions of the distribution and variances of marker effects. rrBLUP assumes that marker effects follow a normal distribution where all effects are shrunk to a similar and small size, while Bayesian methods (BayesA, BayesB, Bayesian LASSO and BayesR) assume that genetic variances specific to the marker effects and including a priori data on the probability distributions of marker effects. The GBLUP method computes the additive genetic merits from a genomic relationship matrix and is equivalent to rrBLUP under conditions that are generally met in practice [14]. The RKHS regression model is a linear combination of the basic function provided by the reproducing kernel [15]. Recent studies have indicated that the selection of suitable statistical methods depends on the actual data at hand and the pattern of phenotypic variation in the traits of interest and with reference population used [9, 16].

Besides statistical methods, other factors are known to influence the accuracy of genomic prediction models, such as the size of the training population, number of markers employed, and relatedness between the training and validation population and, by extension, to the future selection candidates. Hayes et al. [17] found that for a given effective population size (*N*_*e*_), increasing the size of the reference population leads to improved accuracy of GS based predictions. Closer relationship between training population and selection candidates has been reported to lead to a higher accuracy of genomic predictions, while enlarge genetic diversity of the training population resulted in lower accuracy [18]. A number of simulation and empirical studies have shown that increasing the number of markers may improve the predictive accuracy as the *N*_*e*_ also increases [9, 19−21]. However, increasing the number of markers in small *N*_*e*_ populations has little or no improvement on predictive accuracy [22, 23]. Going one step further from previous studies in forest trees, where individuals of the same breeding generation were allocated to training and validation sets for the evaluation of genomic prediction models, in this study we used both the parental and progeny generations of *E. grandis*, *E. urophylla* and their F_1_ hybrids to build prediction models using different subsets of parents and progeny for training and validation. A multi-species single-nucleotide polymorphism (SNP) chip containing 60,904 SNPs [24] were used to provide high-density genotyping of the two generations. Based on these data, we developed genomic prediction models for height, circumference at breast height (CBH), volume, wood basic density and pulp yield, using a number of statistical methods and compared their performance to the traditional pedigree-based prediction. Furthermore, we evaluated the impact of varying the number of SNPs and the training set/validation set composition and size on the predictive ability (PA) of genomic prediction.

## Methods

### Breeding population

The breeding population in this study was established by controlled crossings of 86 *E. urophylla* and 95 *E. grandis* trees (G0 population) following a incomplete diallel mating design, resulting in 16,660 progeny individuals (G1 population) comprising 476 full-sib families with 35 individuals per family. In 2009, the progenies were deployed in a field trial in a randomized complete block design with single-tree plots and 35 reps per family in Belmonte (Brazil, 39.19W, 16.06 S, 210 m above the sea level) at Veracel Celulose S.A. (Eunápolis, BA, Brazil). Our experimental population consists of 168 parents (78 of *E.urophylla* and 90 of *E.grandis*) (G0), as not all parents were still alive at the time of study, and 958 progeny individuals (G1) sampled across 338 full-sib families by avoiding low performing trees. The number of individuals in each full-sib family ranged from one to 13 with an average of 2.8 individuals per family.

### Phenotyping

For the 958 G1 samples, height, volume, and circumference at breast height (CBH) were measured at age three and six years, respectively, and the wood traits (basic density and pulp yield) were measured at age five years. For the 168 G0 parents, the same traits had been measured at age seven years for *E. grandis* and at age five years for *E. urophylla*. Briefly, height was measured using a Suunto hypsometer/height meter (PM-5/1520 series) and CBH was measured with a centimetre tape at 130 cm above ground. Wood properties were estimated by employing near-infrared reflectance spectra of sawdust samples collected at breast height using a FOSS NIRSystem 5000-M and applying calibration models developed earlier by Veracel S.A.

A mixed linear model was applied to minimize the impacts of environmental and age differences on each trait.

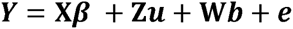

where ***Y*** is a vector of trait; ***β*** is a vector of fixed effects, including overall mean, experimental sites and age differences; ***u*** is a vector of random additive genetic effect of individuals with a normal distribution, 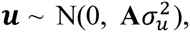 ***A*** is a matrix of additive genetic relationships among individuals; b is a vector of random incomplete block effect nested in each experimental site; and ***e*** is a heterogeneous random residual effect in each experimental **X**, **Z** and **W** are incidence matrices for ***β***, ***u*** and ***b***, respectively. The phenotypes of each trait were then corrected by subtracting variation of sites, ages and blocks effects for all individuals, and are referred to as adjusted phenotypes. The adjusted phenotypic traits were used for calculating the heritability of traits and for building genomic prediction models.

### Genotyping and quality control

The 168 G0 and 958 G1 populations were genotyped using the Illumina Infinium EuCHIP60K [24] that contains probes for 60,904 SNPs. EUChip60K intensity data (.idat files) were obtained through GENESEEK (Lincoln, NE, USA). SNP genotypes were called using GenomeStudio (Illumina Inc., San Diego, CA, USA) following standard genotyping and quality control procedures with no manual editing of clusters as described earlier [24]. Further quality control of the genotyped samples was performed using PLINK [25]. Nine G1 individuals were removed due to low sample call rate (<70%) or high inbreeding coefficient (F>1). 10,240 SNPs were excluded due to low call rate (<70%), 9,243 SNPs were filtered out due to monomorphism or minor allele frequency (MAF) < 0.01, and 117 SNPs were removed due strong deviations from Hardy-Weinberg equilibrium (p-value < 1×10^-6^).

After quality control, missing genotypes of the remaining individuals were filled in by imputation. We first tested the accuracy of imputation methods across a range of missing data (2% – 30%) by artificial removing SNPs from a fraction of our genotypes. Among the available family-based and population based methods we assessed the following programs for imputation accuracy: BEAGLE [26], fastPHASE [27], MENDEL [28], random forest, SVD Impute, k-nearest neighbors [29], BLUP A matrix, Bayesian PCA, NIPALS, Probabilistic PCA [30]. BEAGLE provided the best accuracy for all missing data percentages, with accuracies exceeding 95% in all cases (Additional file 1). We therefore used BEAGLE to impute missing genotypes at the retained 41,304 SNPs across the 168 G0 and 949 G1 individuals. The imputed genotypic data was subsequently used in all genomic prediction analyses. LD between SNP pairs was measured using the squared correlation coefficient (r^2^) for SNPs located on the same chromosome. The decay of LD versus physical distance was then modelled using the nonlinear regression method described in Remington et al. [31].

We further studied the population structure and pairwise genomic relationship among the 1117 individuals by performing principal components analysis (PCA) [32] and kinship analysis [33] using 10,213 independent SNPs (LD-pruned) (r^2^ < 0.2) calculated in PLINK [25]. Pedigree-based genetic relationship was estimated from ABLUP (see below for further information).

### Statistical methods for genomic prediction

Four statistical methods were assessed to estimate the parameters in equation (1) and for predicting GEBVs, including genomic best linear unbiased predictor (GBLUP) [5], ridge regression BLUP (rrBLUP) [6], Bayesian LASSO (BL) [34], and reproducing kernel Hilbert space (RKHS) regression [15]. The performance of the four genomic prediction methods was compared with that of the commonly used pedigree-based BLUP (ABLUP) [35].

The GEBVs were estimated using the following mixed linear model:

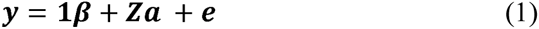

where ***y*** is the vector of adjusted phenotypes of single trait, ***β*** is the vector of overall mean fitted as a fixed effect, ***a*** is the vector of random effects, and ***e*** is the vector of random residual effects. **1** and **Z** are incident matrix of ***β*** and ***a***, respectively.

#### ABLUP

ABLUP is the standard method for predicting breeding values using the expected relatedness among individuals based on pedigree information [35]. For ABLUP, the vector of random additive effects (***a***) in the equation (1) is assumed to follow a normal distribution,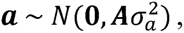 where **A** is the additive numerator relationship matrix estimated from pedigree information and the 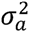 is the additive genetic variance. The residual vector ***e*** is assumed as 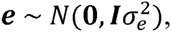 where ***I*** is the identity matrix. Under these assumptions, equation (1) can be re-written as:

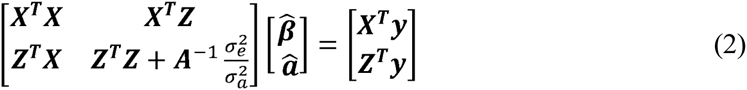

where 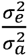 is estimated using a restricted maximum likelihood method. The estimated breeding values (***â***) can be calculated directly from equation (2). ABLUP calculations were performed using ASReml 3.0 [36].

#### GBLUP

The GBLUP method is derived from ABLUP, but differs in that the matrix **A** in equation (2) is replaced with the genomic relationship matrix (**G**) that is calculated from the genotypic data as 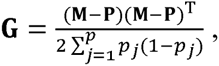 where **M** is the matrix of samples and their corresponding SNPs denoted as 0, 1, 2, **P** is the matrix of allele frequencies with the *j*−^th^ column given by 2(*p*_*j*_− 0.5), where *p*_*j*_ is the observed allele frequency of the samples [5]. In GBLUP, the random additive effects (***a***) in the equation (1) is assumed to follow 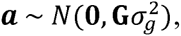 where 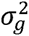 is the genetic variance and GEBVs are again calculated from equation (2) but with ***A−1*** replaced by ***G−1*** and 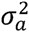 replaced by 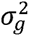 The GBLUP calculations were performed using ASReml 3.0 [36].

#### rrBLUP

As opposed to the previous two methods rrBLUP alters the notations of parameters ***a*** and ***z*** in the equation (1), where ***Z*** now refers to a design matrix for SNP effects, rather than incident matrix and ***a*** refers to SNP effects that are assumed to follow 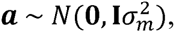 where 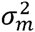 denotes the proportion of the genetic variance contributed by each SNP [6]. With these alterations, equation (2) becomes:

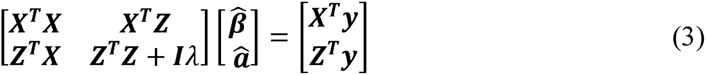

where 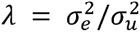 is the ratio between the residual and marker variances. A prediction for the GEBV for each individual is calculated as 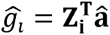 from equation (3), where 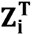 is the SNP vector for individual *i* and **â** is the vector of estimated SNP effects. All calculations were performed using the rrBLUP package in the R environment [33].

#### Bayesian LASSO

The Bayesian LASSO (BL) method is the Bayesian treatment of LASSO regression proposed by Legarra et al. [34]. In BL the vector of SNP effects **a** in equation (1) is assumed to follow a hierarchical prior distribution with 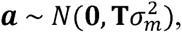 where 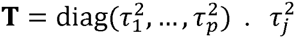 is assigned as 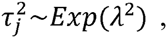 *j*=1,…,*p*. *λ*^2^ is assigned as *λ*^2^ ~*Gamma*(*r*,*δ*). The residual variance 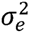 is assigned as 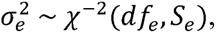

We implemented the BL method using the BLR package in R [37]. Here a Monte Carlo Markov Chains sampler was applied and prior parameters (*df*_*e*_,*S*_*e*_,*r*,*δ*,and *λ*^2^) were defined following the guidelines proposed by de los Campos *et al.* [38]. The chain length was 20,000 iterations, with the first 2,000 excluded as burn-in and with a subsequent thinning interval of 100.

#### RKHS

RKHS assumes that the random additive effects in equation (1) 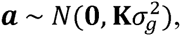 where **K** is computed by means of a Gaussian kernel that is given by K_*ij*_ = *exp*(−*hd*_*ij*_) [15]. *h* is a semi-parameter that controls how fast the prior covariance function declines as genetic distance increase and *d*_*ij*_ is the genetic distance between two samples computed as *d*_*ij*_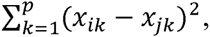 where *x*_*ik*_ and *x*_*jk*_ are *k*th SNPs (*k*=1,…,*p*) for the *i*th and *j*th samples, respectively. We implemented the RKHS method through the BGLR package in R [39], which uses a Gibbs sampler for the Bayesian framework and assigns the prior distribution of 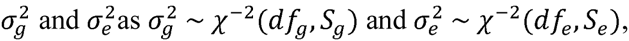 respectively. Here we chose a multikernel model suggested by Perez [39], where three *h* were defined as 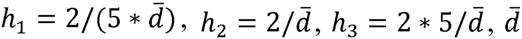 was the median of *d*_*ij*_. The Gibbs chain length was 20,000 iterations with the first 2000 iterations discarded as burn-in and a thinning interval set to 100.

### Heritability estimation

We estimated the pedigree-based narrow-sense heritability 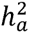 using the relationship matrix from the ABLUP method, and the narrow-sense genomic heritability 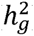 using the genomic relationship matrix from GBLUP [40]. The respective heritabilities were calculated as:

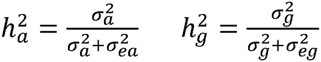

where 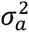 is the additive genetic variance and 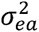 is the residual variance estimated with ABLUP, while 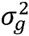 is the genetic variance and 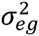 is the residual variance estimated with GBLUP.

### Size and genetic composition of the training and validation sets

We simultaneously assessed the impact of the size and genetic participation of G0 and G1 individuals in the training set (TS) and validation set (VS) of the genomic prediction models. Regarding TS/VS sizes, we divided all 1,117 (G0 and G1) individuals into five different size groups with a ratio of TS to VS of 1:1, 2:1, 3:1, 4:1 and 9:1. The corresponding sizes of the TS/VS were respectively 558/559, 743/374, 836/281, 892/225 and 1003/114. Within these pre-established size compositions, four scenarios of the participation of G0 and G1 individuals were evaluated to assess the impact of varying the degrees of relationship and diversity between TS and VS. In the first scenario (CV_1_) assignment of individuals to either TS or VS was random. For the second scenario (CV_2_) all G0 parents were assigned to the TS and complemented with G1 individuals up to the required number in the set, while the VS was composed exclusively of G1 individuals. The third and fourth scenarios were built based on minimizing and maximizing relatedness between TS and VS. The relatedness-based assignment of individuals was determined using the procedure described in Spindel *et al.* [9]. Briefly, 1,117 individuals were assigned to 182 clusters based on genotypes using the k-means clustering algorithm, a method that attempts to minimize the distance between individuals in a cluster and the centre of that cluster. Using the relatedness estimates, CV_3_ was then built by assigning individuals to TS and VS based on dissimilarity, such that individuals from the same cluster were not allowed to be both in the same TS or VS. For CV_4_ individuals from same cluster were forced to be either in the TS or VS [9].

### Genomic prediction models

We evaluated the effects of the five statistical methods (GBLUP, rrBLUP, BL, RKHS and ABLUP), five TS/VS sizes and four TS/VS composition scenarios (5*5*4 = 100 models in total) on the predictive ability (PA) of genomic prediction. For each of the 100 models, 200 replicate runs were carried out for each trait and the performance of the models were evaluated in terms of their 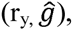, which is defined as the Pearson correlation between the adjusted phenotypes and the GEBVs of the samples in the VS. ANOVA was performed on 80 out of 100 models tested (20 ABLUP models excluded) to partition the variance into different sources, with all effects declared as fixed, comparing all the sources of variation (genomic prediction method, TS/VS size and genetic composition). Significant differences found were further assessed by means of a paired t tests (α = 5%), adjusted by a Bonferroni correction. The 80 models as described above were used for assessing the impact of TS/VS composition and TS/VS size, while all 100 models were used to evaluate the statistical methods against ABLUP. All available SNPs were used in all the analyses of these models.

### Numbers and genomic location of SNPs subsets

We finally assessed the impact of the number of SNPs and their locations (gene vs. intergenic region) on the PA of genomic prediction models. 12 subsets with different numbers of SNPs were generated by randomly selecting 10, 20, 50, 100, 200, 500, 1,000, 2,000, 5,000, 10,000, 20,000 and 41,304 SNPs from all the available SNPs. For SNP location, SNPs subsets located in different regions of the genome were established by including SNPs located in four different regions: (i) coding sequences (CDS) only (11,786 SNPs); (ii) entire genic regions including CDS, UTRs, introns, and sequences 2kb up and downstream of the gene (30,405 SNPs); (iii) intergenic regions (10,899 SNPs), and (iv) all 41,304 SNPs. The location of each SNP was obtained by mapping SNPs onto *E.grandis* genome database using SnpEff [41]. Genomic prediction models were built for all four TS/VS compositions using only the two statistical methods (GBLUP and RKHS) that showed optimal predictive performance in the previous analyses, and the TS/VS size ratio of 4:1 (892/224) were used on the PA evaluations.

## Results

### Phenotypic trait correlations

Growth (height, volume, and CBH) and wood properties (basic density and pulp yield) were measured for all 168 G0 and 949 G1 individuals. The raw phenotypic data were adjusted using a mixed linear model to minimize the impacts of environment and age differences. The pairwise correlations between the adjusted traits were described by calculating Pearson correlation coefficients (Figure 1). Growth traits were correlated with each other. Interestingly, however, while CBH and volume at age three and six years were highly correlated (r = 0.92 and 0.95 respectively), height at age three was only weakly correlated with height at age 6 (r = 0.36). For wood properties traits, basic density was negatively correlated with pulp yield, although weakly (r = -0.28). Growth traits showed no correlations with wood traits (r = - 0.1 to 0.1).

**Figure 1.**
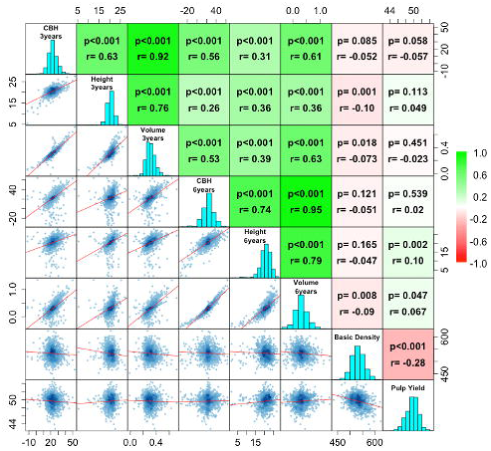
Correlation and distribution of phenotypes. (Scatter plots (lower off-diagonal) and correlations with probability values (upper off-diagonal; H_0_: r=0) for adjusted phenotypes between pairs of traits. Color key on the right indicates the strength of the correlations. Diagonal: histograms of the distribution of adjusted phenotypes values.

### Breeding population structure and relatedness

Population structure across G0 and G1 individuals was assessed by PCA based on 10,213 LD-pruned, independent SNPs (r^2^<0.2). The first two PCs explained 6.07% and 3.8% of the total genetic variance (Figure 2a) and clearly separated the G0 individuals of the two species, *E.grandis* and *E.urophylla,* with the *E.grandis* individuals further subdivided into two subgroups likely representing the two main provenances used in breeding programs in Brazil. The G1 individuals were generally projected into the space defined by their parents, but with a few outliers. The expected pedigree-based and realized genomic-based genomic relationships among G0 and G1 individuals were visualized in heatmaps (blue and red in Figure 2b, respectively). The result of the genomic relationship analysis corroborated the PCA result, in which *E. urophylla* was clustered into a single group, whereas *E. grandis* formed two subgroups. The average values of the realized genomic relationships among what were considered to be full-sibs, half-sibs and unrelated individuals from the pedigree data were generally lower than the expected relationships values (0.309 vs. 0.5, 0.131 vs. 0.25 and .0056 vs. 0, respectively) (Table 1). This result suggests that pedigree errors were likely present in this population. These putative pedigree errors in turn negatively affected the pedigree-based trait heritability, which were considerably lower than those estimated using genomic-based realized genomic relationships (Table 2).

**Figure 2.**
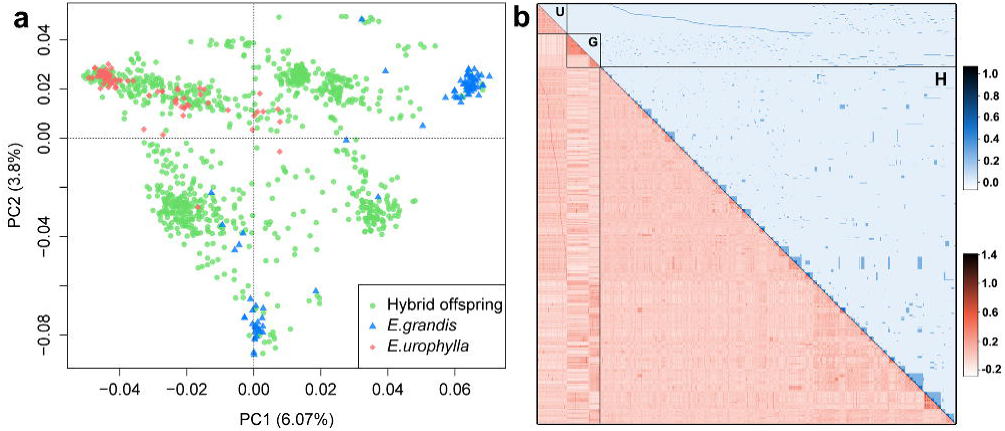
Genetic structure and relatedness in the breeding population. (a) First two principal components of a PCA revealing population structure. Dots represent *E.grandis* (blue), *E.urophylla* (red) and their F_1_ (green) individuals. (b) Heatmaps of the pairwise pedigree-expected relationships (blue, upper off-diagonal) and genomic-realized relationship (red, lower off-diagonal) of the 1117 individuals assigned to *E.grandis* (G), *E.urophylla* (U) and their hybrid progenies (H).

**Table 1.** Pairwise expected pedigree-based and realized genomic-based relationships in the different family types.

|  | Full-sib families<br>(961)<sup>a</sup> |  |  | Half-sib families<br>(12718) |  |  | Unrelated individuals<br>(434252) |  |  |
| --- | --- | --- | --- | --- | --- | --- | --- | --- | --- |
|  | Min | Mean | Max | Min | Mean | Max | Min | Mean | Max |
| Pedigree-expected relationship | 0.5 | 0.5 | 0.5 | 0.25 | 0.25 | 0.25 | 0 | 0 | 0 |
| Genomic-realized relationship | -0.274 | 0.309 | 0.933 | -0.464 | 0.131 | 0.908 | -0.467 | -0.056 | 0.891 |

<sup>a</sup>Number in parentheses indicate the number of pairwise estimates

**Table 2.** Pedigree-based and genomic heritabilities for each trait

|  | CBH (3)<sup>a</sup> | Height (3) | Volume (3) | CBH (6) | Height (6) | Volume (6) | Basic density | Pulp yield |
| --- | --- | --- | --- | --- | --- | --- | --- | --- |
| $h_a^2$<sup>b</sup> | 0.051(0.03) | 0.074(0.04) | 0.057(0.03) | 0.085(0.04) | 0.097(0.05) | 0.068(0.04) | 0.23(0.04) | 0.27(0.05) |
| $h_g^2$ | 0.113(0.04) | 0.171(0.05) | 0.162(0.04) | 0.184(0.04) | 0.193(0.05) | 0.196(0.04) | 0.35(0.05) | 0.46(0.05) |

<sup>a</sup> Number in parentheses correspond the age at measurement;
<sup>b</sup> $h_a^2$ and $h_g^2$ correspond to the pedigree and genomic narrow-sense heritability, respectively, with their standard deviation in parenthesis.

### Predictive abilities with different statistical methods

Estimates of PAs were obtained using different statistical methods, compositions and sizes of TS/VS for each trait (Additional file 2). An ANOVA showed that all these factors had a significant effect on the PA (P-value < 0.005) (Additional file 3). Across the four genomic prediction methods used (GBLUP, rrBLUP, BL, and RKHS) the average PA varied from 0.27 to 0.274 (Additional file 4). All the four methods outperformed the pedigree-based ABLUP prediction (mean PA = 0.121) by an average of 80%-200% across the eight traits (Figure 3). RKHS yielded a slightly better PAs for six out of eight traits and this method was particularly suitable for predicting traits that displayed a lower heritability such as CBH and height. The other three methods generally gave similar results across all traits, although with a slightly better performance than RKHS for pulp yield (Figure 3).

**Figure 3.**
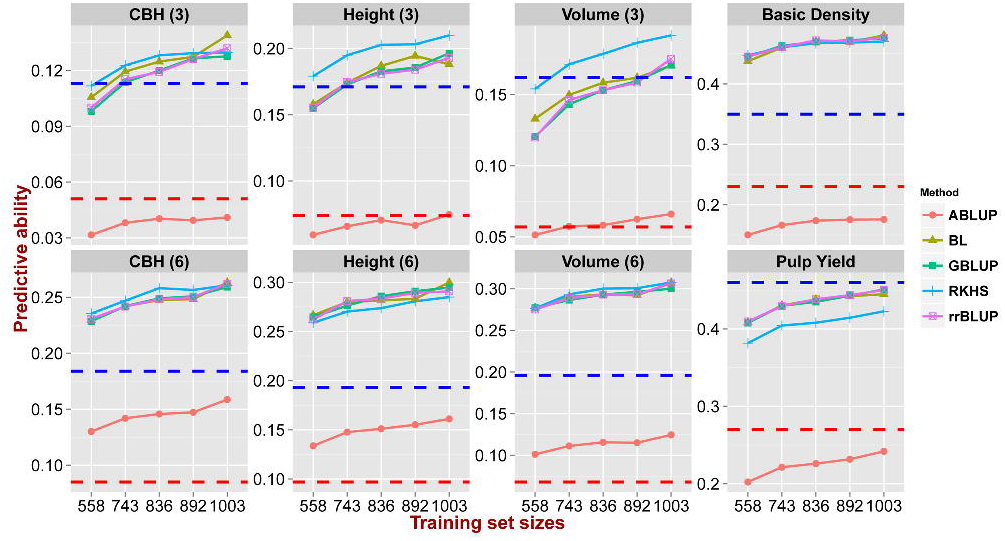
Predictive abilities with different methods and increasing sizes of training sets. Predictive ability (y axis) estimated using five methods across five training set/validation set sizes in numbers of individuals (x axis) 558/559, 743/374, 836/281, 892/225 and 1003/114. Red and blue dashed lines indicate the pedigree-based 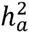 and genomic-realized 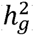 narrow-sense heritability respectively.

### Impact of TS/VS compositions and relative sizes on predictive ability

The average PAs differed significantly for the different TS/VS composition tested varying from 0.253 to 0.286 (Additional file 5). The genomic prediction model built with CV_2_ (all G0 parents in the TS) showed the highest PAs for all traits except pulp yield, whereas models based on CV_3_ (minimum relatedness between TS and VS) gave the worst predictions. The models based on CV_1_ (random assignment) and CV_4_ (maximum relatedness between TS and VS) showed no significant differences in PA (Figure 4, Additional file 5). The average PA was significantly improved from 0.251 to 0.285, as the TS/VS ratio increased from 1:1 (558/559) to 9:1 (1003/113) (Additional file 6), irrespective of the prediction method (Figure 3) or the genetic composition of TS/VS used (Figure 4), clearly showing the importance of an adequate size of the training set used to build prediction models. Furthermore, there was a steeper increase in PA when TS/VS ratio increased from 1:1 (558/559) to 2:1 (743/374) than from 2:1 (743/374) to 9:1 (1003/114) for all traits (Figure 3 and 4).

**Figure 4.**
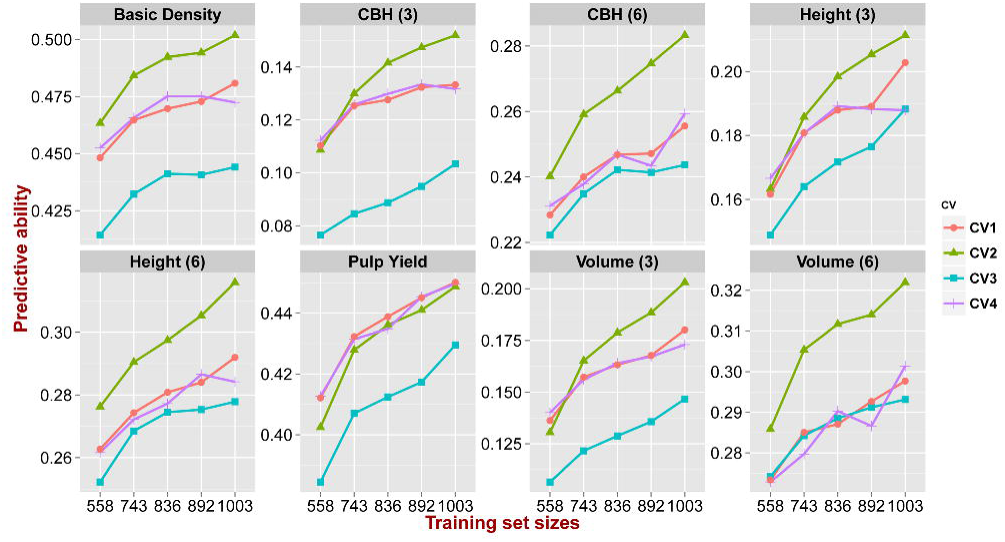
Predictive abilities with variable levels of relatedness between training and validation sets. CV_1_: random assignment of individuals to either training set (TS) or validation set (VS); CV_2_: all the G0 pure species parents assigned to the TS; CV_3_: minimum relatedness between TS and VS individuals; CV_4_: maximum relatedness between TS and VS individuals. Estimates were obtained using GBLUP and RKHS across five TS/VS sizes in numbers of individuals (x axis): 558/559, 743/374, 836/281, 892/225 and 1003/114.

### Impact of the number of SNPs and their genomic location on predictive ability

Estimates of PA using different numbers of SNPs (Additional file 7) and sets of SNPs in different genomic locations (Additional file 8) were obtained with two prediction methods for all the different TS/VS compositions. An ANOVA showed that both the number of genotyped SNPs and their genomic location significantly affect the PA for both prediction methods (GBLUP and RKHS) (P-value < 0.005), and that the number of SNPs has a larger impact than their genomic location (Additional file 9). The average PAs across all traits decreased from 0.278 to 0.113 when the number of SNPs used in the prediction models dropped from 41,304 to only 10, and the reduction was especially strong when the number of SNPs went below 5,000 (Additional file 10). On the other hand, no significant improvement was generally seen in the average of PA when more than 5,000 SNPs were used (Additional file 10, Figure 5). The results obtained for the different traits suggest that traits with lower heritability are more sensitive to the reduction in the number of SNPs (Figure 5). For instance, PA for basic density (h^2^ = 0.35) went from 0.47 to 0.24 (a 50% decrease) when the number of SNPs dropped from 40,000 to 10, whereas CBH of age three (h^2^ = 0.113) decreased from 0.128 to 0.03 (a 77% decrease). Overall, few and only slight significant differences were seen in PAs by using SNP sets located in different genomic regions (Figure 6), the average PAs range from 0.270 to 0.284 (Additional file 11). Predictions using SNPs located in intergenic regions were marginally better than using SNPs in genic regions or all SNPs, except for pulp yield that could be better predicted based on models using SNPs from coding and gene regions (Figure 6). When comparing the PA of models using SNPs in coding versus entire gene regions, the latter had a slightly better performance, most likely due to the larger number of SNPs used (30,504 vs. 11,786) and not to any specific effect of genomic location. When we assessed the pairwise LD (r^2^) amongst the SNPs in the four regions tested, the extent of LD differed among them, with LD showing the most rapid decay in coding regions and the slowest one in intergenic regions (Additional file 12).

**Figure 5.**
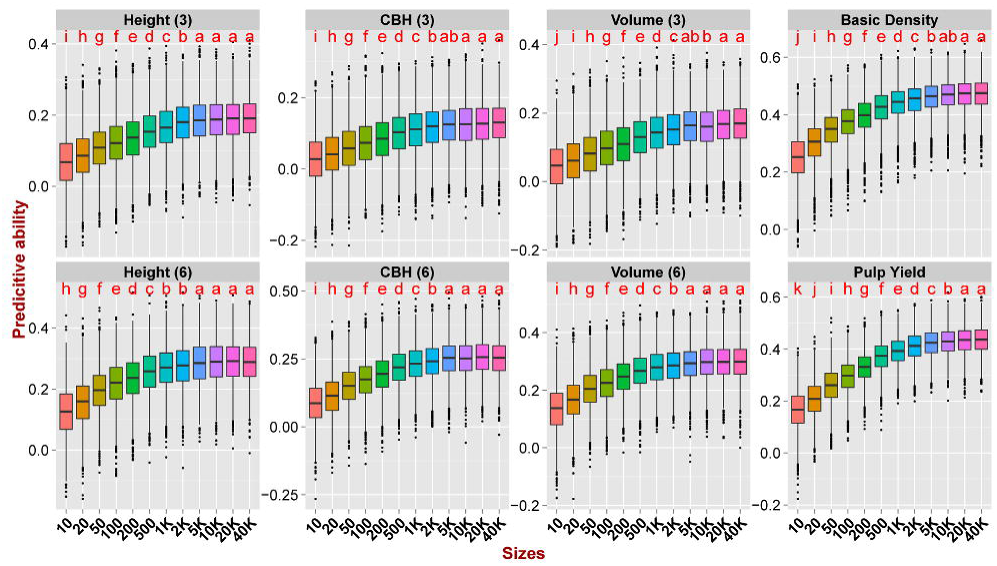
Predictive abilities with increasing numbers of SNPs. Predictive ability estimated with GBLUP and RKHS with increasingly larger sets of SNP sampled at random from the total of 41,304 SNPs. Outliers are indicated by black dots. Letters indicate significant difference between the different models after Bonferroni adjustment (P < 0.05).

**Figure 6.**
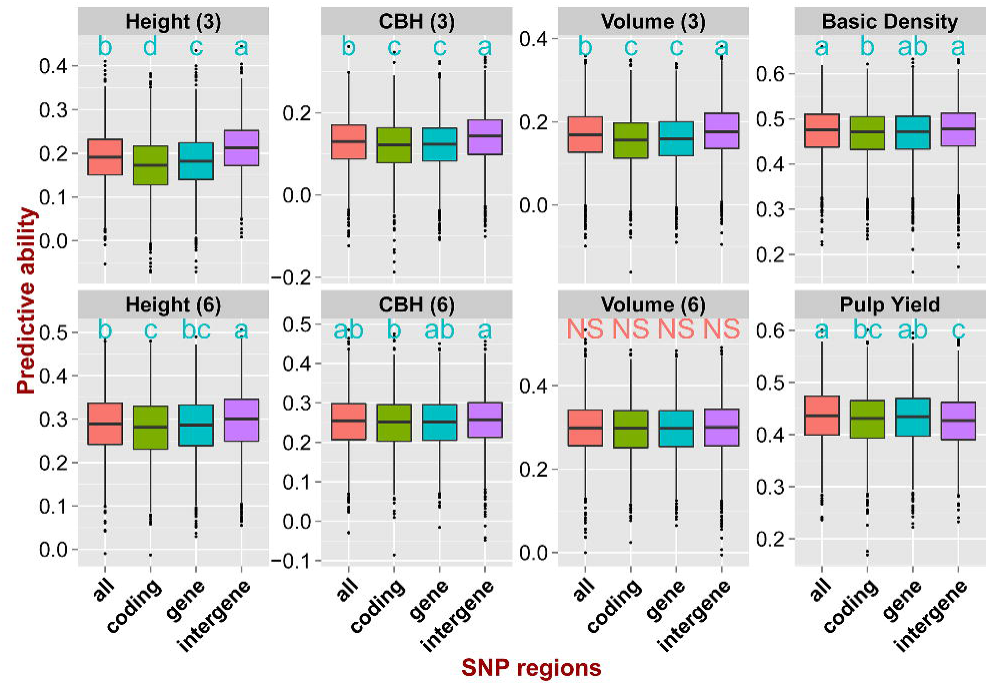
Predictive abilities using SNPs located in different genomic regions. Predictive abilities using SNPs located in different genomic regions. Predictive ability estimated with GBLUP and RKHS using 11,786 SNPs in coding DNA, 30,405 SNPs in genic regions (CDS, UTR, intron, and within 2kb upstream and downstream of genes), 10,899 SNPs in intergenic regions and all 41,304 SNPs. Letters indicate significant difference between the different models after Bonferroni adjustment (P < 0.05).

## Discussion

This study presents the results of an empirical evaluation of the accuracy of genomic prediction of growth and wood quality traits in *Eucalyptus* using data from a high-density SNP array. Our results are based on data from a two generations breeding population and provide additional encouraging results on the prospects of using genomic prediction to accelerate breeding. We have assessed a range of factors, including the statistical methods used to estimate predictive ability, the size and composition of the training and validation sets as well as the number and genomic locations of SNPs used in the prediction model. Hereafter we will discuss how these factors influenced the prediction accuracy.

### Genomic data corrected pedigree inconsistencies

All four genomic prediction methods performed significantly better than the pedigree-based evaluations for all complex traits assessed (Figure 3). While similar results have been reported for animals [16, 42] and crop species [9, 35] across a number of traits, in forest trees prediction accuracies using genomic data have generally been similar or up to 10-30% lower than accuracies obtained using pedigree-estimated breeding values, including *Eucalyptus* [4], loblolly pine (*Pinus taeda*) [43], white spruce (*Picea glauca*) [44, 45], interior spruce (*Picea engelmannii* × *glauca*) [46, 47] and maritime pine (*Pinus pinaster*) [48]. Genomic predictions with lower accuracies than pedigree-based predictions could arise from insufficient marker density, such that not all casual variants are captured in the genomic estimate [40], or an overestimate of the pedigree-based prediction due to its inability of ascertaining the true genetic relationships in half-sib families [46]. Our result however differ from previous studies in forest trees due to the fact that the average pairwise estimates of genetic relationship among individuals were substantially lower using SNP data than expectations based on pedigree information (Table 1), clearly suggesting that the expected pedigrees, and consequently the pairwise relationships, had considerable inconsistencies that were corrected by the SNP data. We speculate that these inconsistencies likely derived from pollen contamination and mislabelling in the process of generating the full and half-sib families. Besides correcting potential pedigree errors, the relatively dense SNP data used in our study also was able to accurately capture the Mendelian sampling variation within families so that genetic variances estimates were based on the actual proportion of the genome that is identity by descent (IBD) or state (IBS) among half- or full-sib individuals, resulting in improved estimates of trait heritability (Table 2).

### Genomic predictions show that traits adequately fit the infinitesimal model

Overall, the different genomic prediction methods provided similar results for the traits evaluated with only a slight advantage for RKHS showing better PAs for growth traits that had lower heritability (Figure 3) although for pulp yield, RKHS instead was the worst performing method. It is possible that the definition of a kernel simply was not suitable for this particular trait [15]. Our results corroborate previous reports both in crops and animals [16, 49, 50], as well as in forest tree studies. In loblolly pine, for example, the performance of rrBLUP and three Bayesian methods was only marginally different when compared across 17 traits with distinct heritabilities, with a small improvement using BayesA only for fusiform rust resistance where loci of relatively larger effect have been described [43]. Similar results were obtained for growth and wood traits in other forest trees studies showing no performance difference between rrBLUP and Bayesian methods [45, 47, 48]. This occurs despite simulation studies suggesting that Bayesian methods, like BL, should outperform univariate methods such as rrBLUP and GBLUP [6, 51, 52]. One possible reason for the apparent disagreement between simulations and empirical data sets is that the true QTL effects for most of traits are relatively small and the distribution is less extreme than simulated data [53]. Our results therefore support the proposal that either rrBLUP or GBLUP are effective methods in providing the best compromise between computation time and prediction efficiency [54] and that the quantitative traits assessed in our study adequately fit the assumption of the infinitesimal model.

### Training set size, composition and relatedness strongly affect predictive ability

Our results show that the size and the variable TS/VS compositions in terms of relatedness between training and validation sets had the largest impact on the PA irrespective of the analytical method used (Figure 4). The average PA rapidly increased with increasing sizes of the TS and did not show any sign of plateauing. Earlier simulations of *Eucalyptus* breeding scenarios had in fact shown that with up to N= 1,000 individuals in the TS, the accuracy would rapidly increase, and additional gains would be seen up to N= 2,000 individuals for lower heritability traits, larger numbers of QTLs involved and larger effective population size (*N*_*e*_). After N= 2,000 the predictive accuracy would tend to plateau irrespective of the *N*_*e*_ and genotyping density [20]. Later simulations mirroring a eucalypt breeding scheme also showed a considerable improvement of genomic predictions with increasing training population sizes by consolidating phenotypic and genotypic data of individuals from previous breeding cycles [55]. Simulations [19, 56] and proof-of-concept studies [57] in crop species also show improved PA with larger TS sizes. Larger training populations alleviate the probability of losing rare favourable alleles from the breeding population as generations of selection advance. Additionally by sampling more individuals for training, a larger diversity is captured and better estimates of the marker effects are obtained which in turn positively impact predictions in cross-validations and future genomic selection candidates.

As expected, relatedness between TS and VS had a large impact on PAs for all traits. Prediction models built under scenario CV_3_ (minimized relatedness between TS and VS) resulted in significantly worse predictions than in scenario CV_4_ when relatedness was maximized. Increasing the genetic relationships between training and selection candidates effectively has the same consequence as reducing the *N*_*e*_ such that the stronger the relationship, the higher in the predictive accuracy. Our results are in line with previous reports in forest trees showing that models developed for one population had limited or no ability of predicting phenotypes in an unrelated one in white spruce [44, 45] and *Eucalyptus* [4], indicating that prediction models will be population specific. With lower relationship between TS and VS, the extent of LD is shorter and not stable across distantly related populations and the predictive ability of genomic prediction model is reduced. Recent simulations show that the accuracy of genomic prediction models decline approximately linearly with increasing genetic distance between training and prediction populations [58]. Increased relatedness reduce the number of independently segregating chromosome segments and therefore increase the probability that chromosome segments IBD sampled in the training population are also found in the selection candidates. Our results provide additional experimental evidence that for successful implementation of GS the selection candidates have to show a close genetic relationship to the training population.

PAs were considerably higher when all the G0 parents were kept in the TS (scenario CV_2_). This result could be due to two reasons. On one hand, by keeping all G0 parents in TS, we had a large diversity available for training, which could explain the positive impact of G0 inclusion on predictions. On the other hand, it is possible that by allocating all G0 individuals to the TS the positive effect we observe could strictly not be due to increased predictive power of including G0 individuals but rather a way to avoid the potentially negative impact of having pure species parents in the validation set in combination with G1 progeny that were largely F_1_ hybrids. In order to evaluate this, we estimated PA of genomic prediction models by using GBLUP and RKHS, having only the 168 G0 parents for TS and randomly selected 168 G1 individuals in VS. To control for the effect of the strongly reduced TS size, we compared this setup with random assignment of individuals to TS or VS but keeping the size of each at N=168. The results showed considerably lower PAs (even zero or negative) when using only pure species parents to predict G1 hybrid progeny phenotypes (Additional file 13). This observation, together with the fact that PAs with scenario CV_4_ (maximum relatedness between TS and VS) were also generally lower than CV_2_, suggesting that the higher PAs we observe for scenario CV_2_ is mostly due to avoiding the negative effect of having pure species parents in the VS.

The issue of genomic prediction in hybrid breeding has been investigated so far only within species and only for domestic animals, more specifically for bovine and pig breeding in which selection is carried out in pure breeds but the aim is to improve crossbred performance [42, 59]. Results from simulations show that training on crossbred data provides good PAs by selecting purebred individuals for crossbred performance, although PAs drop with increasing distances between breeds [60]. When crossbred data is not available, separate purebred training populations can be used either separately or combined depending on the correlation of LD phase between the pure lines [61], which in turn is in part determined by the time of divergence between the populations. Compared to bovine breeds that belong to the same species and have diverged relatively recently (<300KYA) [62], the estimated divergence time between the two *Eucalyptus* species used in our study is much older, estimated at 2-5 MYA [63]. Therefore, no correlation of LD phase between these two species is expected and it is not surprising that training on the combined pure species sets to validate on the F_1_ hybrids resulted in poor PA. To the best of our knowledge, our results are the first ones to provide an initial look at the issue of genomic prediction from pure species to interspecific hybrids indicating that, consistent with expectations, models have to be trained in hybrids if one is to predict phenotypes in hybrid selection candidates.

### Number of SNPs is more important than SNP genomic location

Across all traits, no major improvement was detected in PA when more than 5,000 SNPs were used (Additional file 10, Figure 5), although a slight increase were observed for height of age three, basic density and pulp yield when using GBLUP based on 20,000 SNPs. Several studies have also shown that considerably lower numbers of SNPs provided PAs equivalent to those observed using all SNPs available [22, 64]. The necessary number of SNPs needed for genomic prediction model depends on the extent of LD, which strictly related to *N*_*e*_. Our results, where we achieve equivalent PAs using either all of 12–20% of the genotyped markers suggests that it represents a closed breeding population with a relatively limited *N*_*e*_. This has been a common approach in domestic animals with the intent of developing low-density genotyping chips to reduce genotyping costs [8]. The main advantage of using reduced SNP panels is cost-effectiveness, although it is expected that using a higher density of markers will be necessary to mitigate the decay of PAs over generations due to the combined effect of recombination and selection on the patterns of LD [65]. It is also questionable whether it will be more cost effective to have targeted low-density SNP chips for specific populations or a full SNP chip that can be used across breeding populations of several organizations. By having a SNP chip that will accommodate several populations the cost-effectiveness and economy of scale of amassing many more samples to be genotyped with the same chip will likely be much larger than the cost reduction observed by using a smaller number of SNPs on each specific population.

SNP location also contributed to the predictive ability of genomic prediction model although the effects were rather modest. PAs using SNPs in intergenic regions were slightly better than using SNPs in genic regions or using all SNPs, except for pulp yield that could be somewhat better predicted with SNPs in coding and gene regions (Figure 6). This likely represents a random sampling effect and not any specific enrichment for functional variants for this trait. However, the decline of LD was slower for SNPs in intergenic regions when compared to SNPs in gene and coding regions (Additional file 12) and the slightly longer range of LD might help explain why using SNPs in intergenic regions provided better PAs. With slower LD decay, SNPs in intergenic regions might better capture QTLs across longer genomic segments than SNPs in coding regions where LD decays more rapidly.

## Conclusions

Our experimental results provide further promising perspectives for the implementation of genomic prediction in *Eucalyptus* breeding programs. Genomic predictions largely outperformed the pedigree-based ones in our experiment, mainly due to the fact that our expected pedigree had major inconsistencies, such that all pedigree-based estimates were grossly underestimated. This unexpected result illustrated an additional advantage of using SNP data and genomic prediction in breeding programs. While the main advantage of genomic prediction in eucalypt breeding will likely be the reduction of the breeding cycle length [4], the use of a genomic relationship matrix allowed us to obtain precise estimates of genetic relationship and heritability that we would otherwise not have had access to. Furthermore our results corroborated the key role of relatedness as a driver of PA, the potential of using lower density SNP panels, and the fact that growth and wood traits adequately fit the infinitesimal model such that GBLUP or rrBLUP represent a good compromise between computation time and prediction efficiency. In contrast to previous studies in *Eucalyptus*, we had access to both the pure species parents (*E. grandis* and *E. urophylla*) and their F_1_ progeny. We show that models trained on pure species parents do not allow for accurate prediction in F_1_ hybrids, likely due to the strong genetic divergence between the two species and lack of consistent patterns of LD between the two species and their hybrids.

Several issues remain to be investigated for the operational adoption of genomic prediction in eucalypt breeding. First, how does the accuracy of genomic prediction decline over successive generations of selection due to subsequent recombination? Second, how stable are genomic prediction models across multiple environments and how important is it to consider genotype by environment interactions in the models? Finally, we have only considered additive genetic variance for building genomic prediction models in our population, but it is possible and perhaps even likely that non-additive genetic effects will play an important role in many breeding populations and specifically in populations consisting of early generation hybrids.

## Declarations

### Ethics approval and consent to participate

Not applicable

### Consent for publication

Not applicable

### Data availability

The data that support the findings of this study are available from Veracel but restrictions apply to the availability of these data, which were used under license for the current study, and so are not publicly available. Data are available from the authors upon reasonable request and with permission of Veracel.

### Authors’ contributions

BT, BS and PKI conceived and designed the experiment; GSM phenotyped data; GSM and KZF collected samples for genotyping; DG prepared the DNA for genotyping; BT analysed the data and drafted the first version of the manuscript; DG and PKI provided guidance during data analyses; BT, DG, BS and PKI critically contributed to the final version of the manuscript. All authors read and approved the final manuscript.

## Acknowledgements

We would like to thank Michelle Bayerl Fernandes for her contribution on phenotyping the breeding population. The computations were performed on resources provided by the Swedish National Infrastructure for Computing (SNIC) at UPPMAX and HPC2N.

## List of abbreviations

BL: Bayesian LASSO
CBH: circumference at breast height
CDS: coding sequences
GBLUP: genomic best linear unbiased predictor
GEBV: genomic estimated breeding values
GRM: genomic relationship matrix
GS: genomic selection
IBD: identity by descent
IBS: identity by state
LD: linkage disequilibrium
MAS: marker-assisted selection
*N*_*e*_: effective population size
PA: predictive ability
PCA: principal components analysis
QTLs: quantitative trait loci
RKHS: reproducing kernel Hilbert space
rrBLUP: ridge-regression best linear unbiased prediction
SNP: single-nucleotide polymorphism
TS: training set
VS: validation set

## Additional files

### Additional file 1

Average accuracy of SNP imputation methods with increasing proportions of missing data. SNPs on chromosomes 6 and 8 were randomly removed from the dataset to generate specific missing data proportions. Accuracy between imputed and true SNP genotypes were subsequently calculated with the different methods. (DOCX 1.8Mb)

### Additional file 2

Predictive abilities on genomic selection model that comprises of statistical methods, genetic compositions and relative sizes of Training Set/Validation Set for each trait. (XLSX 17 kb)

### Additional file 3

ANOVA analysis of sources of variation affecting the predictive ability. (DOCX 50 kb)

### Additional file 4

Mean and standard deviation of predictive ability with the five prediction methods for the eight traits. (DOCX 99kb)

### Additional file 5

Mean and standard deviation of predictive ability estimated with the four Training Set/Validation Set compositions. (DOCX 87kb)

### Additional file 6

Mean and standard deviation of predictive ability estimated with the five relative sizes of Training Set/Validation Set expressed in proportions and numbers of individuals. (DOCX 91kb)

### Additional file 7

Mean and standard deviation of predictive ability across increasing numbers of SNPs, statistical methods (RKHS and GBLUP), four Training Set/Validation Set compositions for each of eight traits (XLSX 62kb)

### Additional file 8

Mean and standard deviation of predictive ability estimated with SNPs in four genomic locations, with two statistical methods (RKHS and GBLUP), four Training Set/Validation Set compositions for each of eight traits (XLSX 59kb)

### Additional file 9

ANOVA of predictive ability with SNP genomic location and SNP number as sources of variation. (DOCX 63kb)

### Additional file 10

Average predictive ability estimated with different numbers of SNPs fitted into the model. (DOCX 138kb)

### Additional file 11

Average predictive abilities estimated using SNP sets located in different genomic regions. (DOCX 83kb)

### Additional file 12

Decay of linkage disequilibrium (LD) with physical distance estimated with SNPs in different genomic locations. (a) A comparison of the decay of LD with physical distance in four classes of SNPs located with coding, genic, intergenic and all regions, respectively. Dots of pairwise LD versus physical distance and the LD decay for SNPs located in all regions (b), coding region (c), genic region (d) and intergenic region (e), respectively. (DOCX 1.4Mb)

### Additional file 13

Predictive abilities by training in pure species eucalypt parents and predicting in their F_1_ hybrids. Predictive ability estimated under three training/validation sets (TS/VS) scenarios with two methods (GBLUP and RKHS) for each trait. PO168 (red boxes): all 168 *E. grandis* and *E. urophylla* pure species G0 parents used for training and 168 G1 random selected hybrid progeny for validation; random168 (green): randomly selected 168 individuals from all 1,117 for TS and 168 randomly also for VS; random558 (blue): randomly divided all 1,117 individuals into TS and VS of same size (558/558). Outlier estimates are indicated by black dots. (DOCX 179kb)

